# Human subcutaneous and visceral adipocyte atlases uncover classical and specialized adipocytes and depot-specific patterns

**DOI:** 10.1101/2023.09.04.555678

**Authors:** Or Lazarescu, Maya Ziv-Agam, Yulia Haim, Idan Hekselman, Juman Jubran, Ariel Shneyour, Danny Kitsberg, Liron Levin, Idit F Liberty, Uri Yoel, Oleg Dukhno, Miriam Adam, Antje Körner, Rinki Murphy, Matthias Blüher, Naomi Habib, Assaf Rudich, Esti Yeger-Lotem

## Abstract

Human adipose depots are functionally distinct. Yet, recent single-nucleus RNA-sequencing (snRNA-seq) analyses largely uncovered overlapping/similar cell-type landscapes. We hypothesized that adipocytes subtypes, differentiation trajectories, and/or intercellular communication patterns could illuminate this depot similarity-difference gap. For this, we performed snRNA-seq of human subcutaneous and visceral adipose tissue. Whereas the majority of adipocytes in both depots were ‘classical’, namely enriched in lipid metabolism pathways, we also observed ‘specialized’ adipocyte subtypes that were enriched in immune-related, extracellular matrix deposition (fibrosis), vascularization/angiogenesis, or ribosomal processes. Pseudo-temporal analysis suggested a developmental trajectory from adipose progenitor cells to classical adipocytes via specialized adipocytes, suggesting that the classical state stems from loss, rather than gain, of specialized functions. Lastly, intercellular communication routes were consistent with the different inflammatory tone of the two depots. Jointly, these findings provide a high-resolution view into the contribution of cellular composition, differentiation, and intercellular communication patterns to human fat depot differences.

## INTRODUCTION

Human white adipose tissue is spread anatomically throughout the human body. The largest mass is the human subcutaneous adipose tissue (hSAT) that provides mechanical, chemical and thermal isolation between the inner-body and outer environments. Other major depots surround organs either in the retroperitoneum, or in the body cavities - chest, peritoneal and pelvic. Such organs – the viscera – give these adipose tissues their designation as (human) visceral adipose tissues (hVAT). Classically, VAT provide mechanical and thermal protection to the viscera and retroperitoneal organs. Yet, the past ~30 years revolutionized our understanding of adipose tissues and their roles in human health and disease. All adipose tissues (at their various anatomical locations) were found to be highly secretory, engaging through peptides, lipids, nucleotides and exosomes in active auto-paracrine and endocrine communication, through which they regulate fundamental processes including energy metabolism, immune function, vascular tone and reproduction.

Anatomically, hSAT and hVAT differ at multiple levels. They are well-established to differently impinge on health in pathophysiological states such as obesity, in which excessive hVAT accumulation is much more detrimental to health than preferential hSAT expansion^1–3^. But even in normal physiology the two fat depots differ substantially: Metabolically, VAT is more lipolytic, possibly due to being more sympathetically-innervated^4^; Endocrinologically – the two depots differ in the expression and secretion of adipokines and other secretory product^5^; Immunologically, hVAT tends to assume a more inflamed state, with more pro-inflammatory cell types and polarization states^6^. Collectively, these differences manifest with very distinct transcriptional landscapes when compared head-to-head by whole-tissue/bulk transcriptomic approaches (reviewed in ^7^). Intriguingly, differentially-expressed genes between hVAT and hSAT reveal multiple developmental genes, suggesting even distinct developmental paths of the tissues^8^.

Recently, several human adipose tissue single-cell resolution atlases have been reported^9–15^. Adipocytes, whose variation in size up to 20-fold differences, fragility, and high level of lipids, made them unamenable to single-cell RNA-sequencing, were included in recent atlases that utilized spatial and/or single-nucleus transcriptomics (snRNA-seq). These studies revealed adipocyteś spatial arrangements and differential insulin responsiveness^11^, adipocytes that regulate thermogenesis^13^, plasticity of mouse AT in response to diet-induced obesity^14^, adipogenic potential of distinct progenitor cells^12^, associations between specific adipose tissue cell types and metabolic states^9,10^, as well as commonalities and differences across species and dietary conditions^9^. While depot differences have been noted, these previous human adipose tissue cell atlases mainly emphasized commonalities in cell composition between SAT and VAT, including in adipocyte subtypes. Hence, a major gap in our understanding still remains: Functionally and at the whole-tissue transcriptomics level, hSAT and hVAT are very distinct; by contrast, at the single-cell level, particularly when focusing on the adipocyte subtype landscape – the two fat depots overall seem quite similar.

In the present study, we utilized snRNA-seq of human adipose tissue samples to critically compare adipocytes from hSAT versus hVAT at the single-cell level. We hereby describe depot-specific adipocyte-subtype atlases and highlight commonalities and differences between the depots in: i. adipocyte subtype composition; ii. differentiation trajectories, and iii. inter-cellular communication patterns. These offer a novel, high-resolution view, into the adipocyte-subtype composition, differentiation and function of these depots.

## RESULTS

### Human fat depot-specific adipocyte atlases

To generate hSAT and hVAT depot-specific human adipocyte atlases, we performed single-nucleus RNA sequencing (snRNA-seq) on five paired samples of hSAT and hVAT adipose tissues, and five additional samples of hVAT. Samples were donated by a relatively heathy cohort of men (30%) and women (70%) undergoing elective abdominal surgeries, with diverse BMI (23.8 - 44.1kg/m^2^), age (24.5 – 53.4 years), and backgrounds (Methods, **Figure 1A**). A total of 37,879 hSAT nuclei and 83,731 hVAT nuclei passed quality control scrutiny (as detailed in methods, **Table S1**), and undergone separate clustering based on the nuclear RNA repertoires.

**Figure 1.**
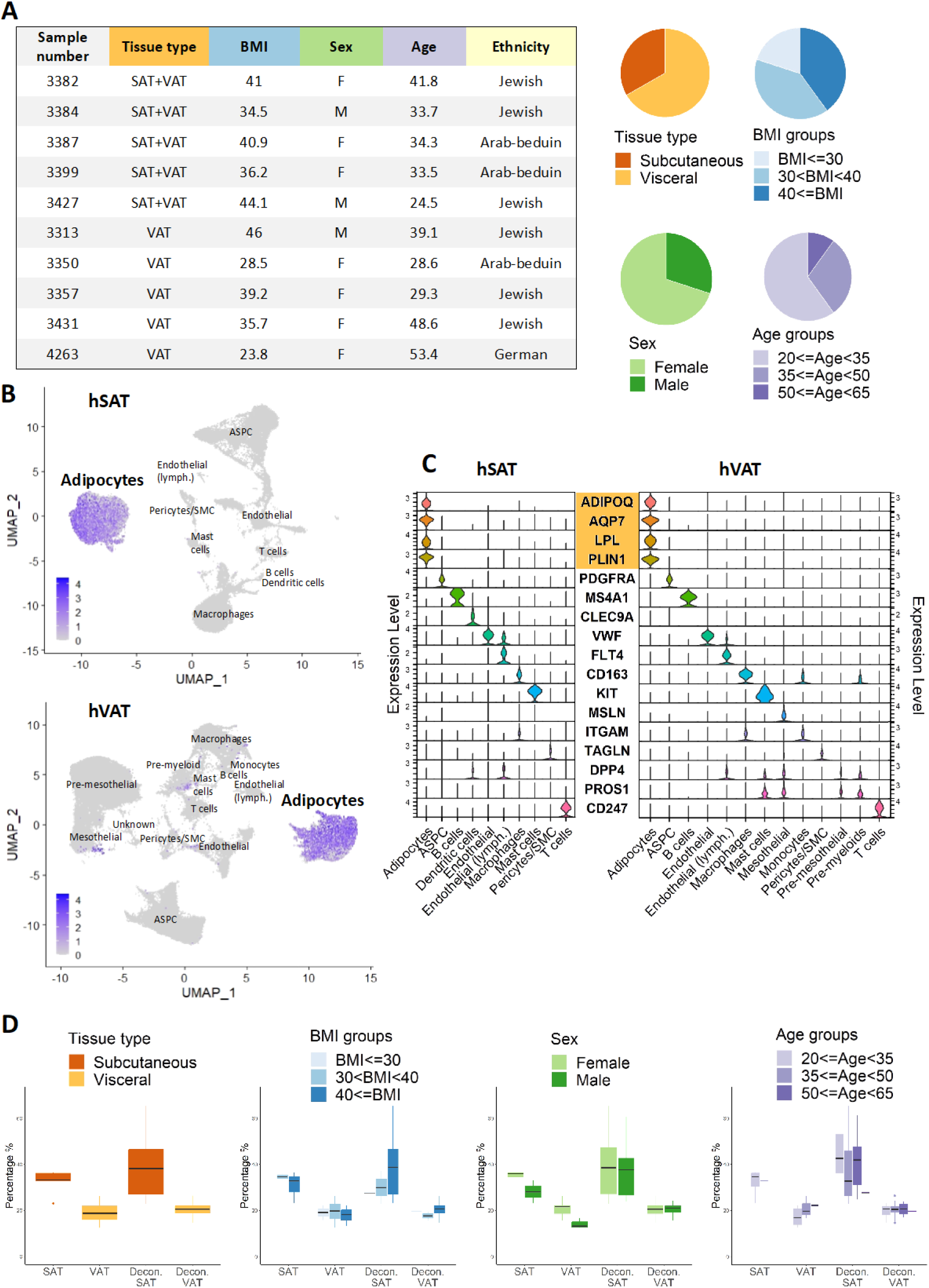
Single-nuclei atlases of human subcutaneous (hSAT) and visceral (hVAT) adipose tissues. A. Clinical parameters of the 15 samples used to build the atlases. Samples were collected from female and male subjects with varying age and BMI, as described in the pie charts. B. The expression of the adipocyte-marker adiponectin (ADIPOQ) in UMAP representations of hSAT and hVAT snNA-seq atlases containing 37,879 and 83,731 nuclei, respectively. In both depots, adipocytes constitute a well-defined cluster. C. The expression of cell-specific markers in different clusters of each depot. Adipocytes were distinguished by multiple markers (marked in orange). D. The fraction of adipocytes per depot, stratified by sex, age, and BMI in the snRNA-seq in-house data (SAT, VAT) and upon estimating their proportions using deconvolution analysis of 73 paired bulk RNA-seq profiles (‘Decon.’ SAT and VAT). Adipocytes were significantly more prevalent in hSAT versus hVAT in both datasets (p<0.001). Other differences, such as between sexes, were statistically significant only in the snRNA-seq data. Statistical significance was assessed using Wald test for snRNA-seq data, and Mann-Whitney for bulk RNA-seq data.

Adipocytes were a readily identifiable and large cluster in both hSAT and hVAT based on expression of adiponectin (*ADIPOQ*) (**Figure 1B**), the cell-type marker used to signify adipocytes in previous studies, as well as the unique expression (compared to other cell-type clusters) of additional adipocyte markers *AQP7*, LPL and PLIN1 (**Figure 1C**). Moreover, genes that were overexpressed (over 2-fold change compared to all other nuclei of that depot, adjusted p-value<0.05, **Table S2**) in clusters identified as adipocytes, were enriched for multiple pathways of fatty acid and lipid metabolism, fitting with adipocytic characteristics (**Table S3**).

Overall, 12,205 (32.2%) and 15,460 (18.5% of total) hSAT and hVAT adipocytes, respectively, were identified. We assessed their fraction out of the total adipose tissue nuclei according to the different strata of donors’ characteristics (**Figure 1D, Table S1**). Adipocytes were significantly more abundant in women versus men in both fat depots (**Figure 1D**). Adipocytes were also more abundant in hSAT versus hVAT, potentially due to the lower cellular diversity of hSAT, which contained 10 cell types, versus hVAT, which contained 13 cell types. These results were corroborated upon analyzing our paired hSAT-hVAT samples alone (**Figure S1**). To test them more broadly, we analyzed 73 paired hSAT and hVAT bulk RNA-seq profiles of patients undergoing bariatric surgeries BMI (54.5 ± 9.3 kg/m^2^). The proportions of adipocytes, which we estimated using a deconvolution tool tailored for snRNA-seq of adipose tissue^16^, supported the larger proportion of adipocytes in hSAT compared to hVAT (Methods, **Figure 1D**).

To investigate the diversity of the adipocyte population in a depot-specific manner, we subjected adipocytes to sub-clustering analysis based on their nuclear RNA profile. This analysis revealed seven and eight adipocyte clusters for hSAT and hVAT, respectively, which we denoted SA1-7 and VA1-8 (**Figure 2A**). Each adipocyte cluster contained nuclei from all samples (**Table S4**), and expressed adipocyte-specific markers at similar levels (**Figure 2B**). Additionally, each adipocyte cluster overexpressed at least 20 genes compared to all other adipocyte clusters, although the degree of overexpression varied (**Figure 2C**). For example, in the largest cluster per depot, namely SA1 and VA1 (85% and 75% of the adipocyte nuclei in hSAT and hVAT, respectively), the overexpression of the 20 topmost genes was modest relative to the 20 overexpressed genes of the other clusters (**Figure 2C**). To unbiasedly compare the adipocyte landscape between the two depots, we combined our two depot-specific datasets, re-clustered them into depot-integrated adipocyte clusters, and assessed the composition of each integrated cluster (denoted ‘integrated Adipocytes – Subcutaneous and Visceral’ (iASV), **Figure 2D, Table S5A**). Notably, SA1-5 and VA1-5 contributed to iASV1-5, respectively, demonstrating the similarity in diversity of the adipocyte population of the two depots. This is illustrated by over 98% of SA1 nuclei and over 99% of VA1 nuclei that clustered together into iASV1 (**Figure 2E-F**). Notably, we also observed depot-differences in adipocytes composition, as iASV6 was contributed predominantly by VA6 (68% of iASV6 cells), while iASV7 was the main recipient of SA7 nuclei (84% of SA7).

**Figure 2.**
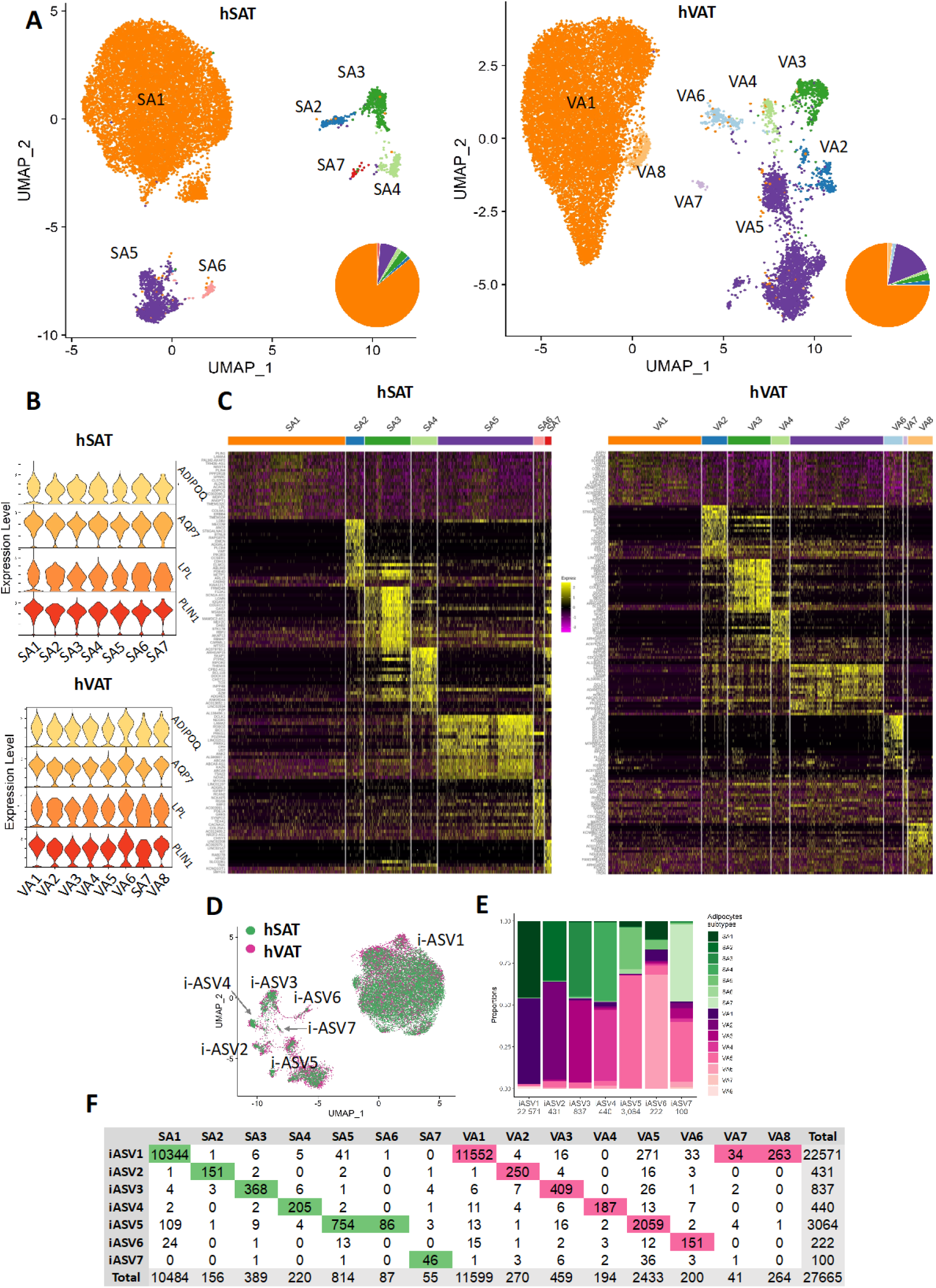
Adipocyte-specific atlases of SAT and VAT. A. UMAP representations of hSAT and hVAT adipocyte atlases containing 12,205 and 15,460 nuclei, respectively. Pie charts show the relative abundance of each adipocyte subset. B. The expression of adipocyte markers in adipocyte clusters of each depot. Distinct clusters expressed markers at similar levels. C. Heatmaps based on the top 20 markers of each adipocyte cluster in hSAT and hVAT. D. UMAP representation of the integrated hSAT and hVAT adipocyte atlases, showing that most subsets overlap between the two depots. E. The proportions of SA1-7 and VA1-8 in each integrated adipocyte SAT-VAT cluster (iASV1-7). The total number of nuclei per integrated cluster appears below the cluster name. F. The number of nuclei from SA1-7 and VA1-8 (columns) that contribute to each integrated cluster (rows). Per column, the entry corresponding to the maximal value (i.e., majority of nuclei) is highlighted.

To gain insight into the putative functional identity of the different adipocyte clusters in each depot, we subjected SA1-7 and VA1-8 separately to gene set enrichment analysis based on their differential expression profiles compared to other adipocyte clusters in the same depot (Methods, **Table S6**). In both fat depots, the large SA1 and VA1 clusters were enriched for fatty acid and lipid metabolic processes, as predictable for adipocytes, and were therefore annotated as “classical adipocytes” (**Figure 3A, Table S6**). These classical adipocytic functions, relatively common to all adipocyte subtypes, explain the modest overexpression pattern observed for these clusters in the differential genes’ heatmap (**Figure 2C**). Enrichment analysis of the other common adipocyte clusters revealed unique and diverse functions that signified specialized adipocyte identities: ‘Angiogenic adipocytes’ (SA2, VA2) were characterized by enrichment for angiogenesis, blood vessel, and vascular development processes (**Figure 3B, Table S6**). ‘Immune-related adipocytes’ (SA3, VA3), were enriched for immune-regulation processes, such as ‘positive regulation of phagocytosis’ and ‘macrophage activation’ **(Figure 3C, Table S6)**. Another subset of immune-related adipocytes (SA4, VA4) was also enriched for lymphocytic processes, including ‘T-cell activation’ and ‘adaptive immune response’, and thus denoted as ‘immune-related (adaptive) adipocytes’ **(Figure 3D, Table S6)**. ‘Extra-cellular-matrix (ECM) adipocytes’ (SA5, VA5), were enriched for ECM processes, including ‘ECM structural constituent’ and ‘ECM organization’ (**Figure 3E, Table S6)**. The depot-specific adipocyte cluster VA6 was enriched in ribosomal components and processes (‘ribosomal subunits’, ‘protein targeting to ER’, **Figure 3F, Table S6),** and thereby termed ‘ribosomal adipocytes’ **(Figure 3F).** SA6 and SA7 in hSAT, and VA7 and VA8 in hVAT, contained small numbers of nuclei and were not significantly enriched for specific processes, and thus remained unannotated. Given that SA2-7 and VA2-8 were not markedly enriched in pathways classically attributed to adipocytes, we jointly relate to these adipocyte subtypes as hSAT and hVAT “specialized adipocytes”, respectively.

**Figure 3.**
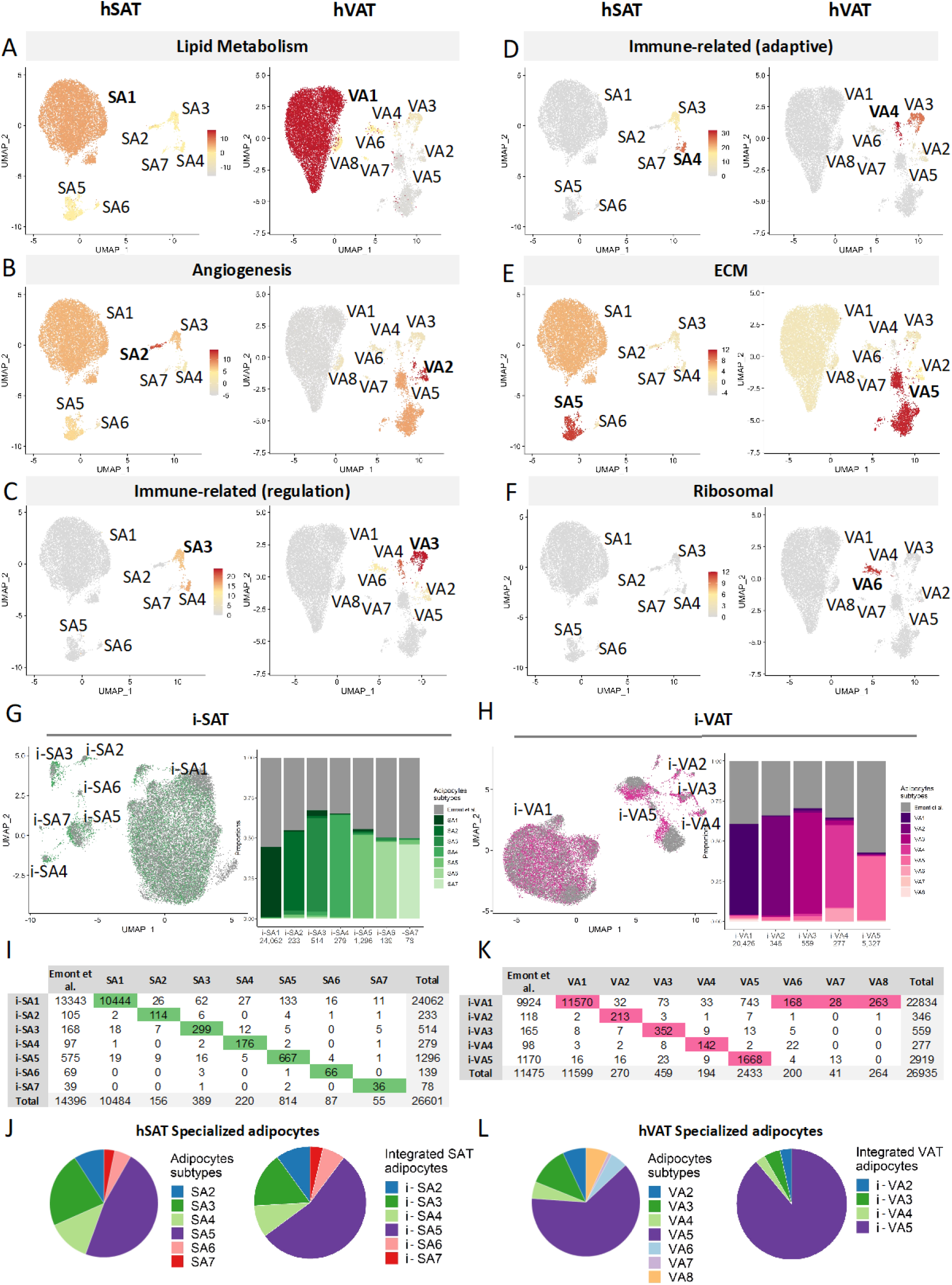
Characterization of adipocyte clusters of SAT and VAT. A-F. UMAP representations of hSAT and hVAT adipocyte atlases. Each cluster was colored by its enrichment for Gene Ontology annotations related to the process specified in the panel. Adipocyte subtypes were enriched for distinct processes, suggesting that they have different functionalities. G-H. UMAP representations of the integration of our in-house adipocyte atlases (green, hSAT; pink, hVAT) and the atlases of Emont et al. (gray). The proportions of SA1-7 and VA1-8 in each integrated SAT (i-SA1-7) or VAT (i-VA1-5) cluster are depicted in bar-plots. The total number of nuclei per integrated cluster appears below the cluster name. I. The number of nuclei from SA1-7 and Emont et al. (columns) that contribute to each integrated cluster (rows). Per column, the entry corresponding to the maximal value (i.e., majority of nuclei) is highlighted. J. The distributions of nuclei among specialized adipocyte clusters for the in-house dataset (SA2-7, left) and for the Emont et al. atlas (i-SA2-7) are similar. K. Same as I, for hVAT. L. Same as J, for hVAT (VA2-8, i-VA2-5).

We next tested whether specialized adipocytes were also present in other atlases of human adipose tissues. Firstly, we integrated our data with 14,396 hSAT and 11,475 hVAT adipocytes from Emont et al.^9^. Integration of the datasets was performed separately for SAT and VAT, using a similar approach to that described above (and further detailed in Methods). Specifically, we re-clustered the adipocyte nuclei in a depot-specific manner, and assessed the composition of each integrated cluster (i-SA in SAT, i-VA in VAT, **Figure 3G-H, Table S5B-C**). In SAT, we identified seven integrated clusters (**Figure 3G**). Although Emont et al., contributed more SAT adipocytes than our in-house data, i-SA1-7 reassuringly matched our in-house clusters SA1-7 (**Figure 3I**). Of the Emont et al., SAT adipocytes, most (92.7%) appeared in the integrated “classical adipocytes” i-SA1 cluster. The remaining 7.3% of the adipocytes segregated into each of the remaining integrated clusters, with each cluster containing 10-100’s of nuclei, and distributed similarly to our in-house data (**Figure 3J, Table S5B**). In VAT, we identified five integrated clusters (i-VA1-5), which matched our in-house clusters VA1-5 (**Figure 3K**; the small VA6-8 were integrated into i-VA1). Of the Emont et al., VAT adipocytes, 86.5% integrated with the “classical” i-VA1 cluster, while the remaining 13.5% split into each of the other specialized adipocytes clusters (**Figure 3L, Table S5C**). Secondly, we applied machine-learning classification methods aimed at classifying adipocytes into our adipocyte subtypes. Specifically, we trained subtype-specific classifiers on our in-house adipocyte subtypes and then applied them to the Emont et al., adipocytes (Methods, **Figure S2A**). This analysis too revealed the presence of considerable fractions of specialized adipocytes per depot (**Figure S2B**, **Table S7**). Lastly, we used deconvolution analysis to estimate the proportions of classical versus specialized adipocytes in the 73 paired hSAT and hVAT bulk RNA-seq profiles (Methods). Classical adipocytes were estimated to be 93.9% of the adipocyte cells in hSAT and 88.5% in hVAT (**Figure S2C**). Jointly, complementary approaches and different datasets support the identification of classical and specialized adipocytes in the two fat depots.

### Differentiation paths towards classical and specialized adipocyte

Adipose tissues harbor a large and diverse population of mesenchymal stem cells with different levels of differentiation potency to a range of cell types (terminal differentiation states). Among them, the sub-population of cells most committed to become mature adipocytes are termed pre-adipocytes, and the entire cell population is characterized by expressing *PDGFRA*, and termed adipose stem and progenitor cells (ASPC in ref.^9^, FAP in ref.^10^). To investigate potential differentiation paths in the two depots and towards classical and specialized adipocyte subtypes, we leverage the single-cell type transcriptome data at hand using trajectory analysis tools^17,18^.

In hSAT, ASPCs were easily identified in a *PDGFRA*-positive cluster, comprising 26% of the entire nuclei population (**Figure 4A**). hSAT-ASPCs could be further divided into five sub-clusters, ASPC1-5 (**Figure 4B**), distinguished by markers that overlapped with previously observed ASPC markers (**Figure 4C**). In particular, of the three largest ASPC clusters (**Figure 4D**), ASPC1 (32% of the ASPC cells) expressed markers of highly proliferative multi-potent progenitor cells^9^, such as DPP4^19^ and CD55^10^, and was F3 negative^19^ (in ref 8 referred to as Adipose stem cells) (**Figure 4C**). ASPC2 (55%) expressed PPARG most highly and was CD9 low, suggesting its commitment to adipocyte fate^20^. ASPC3 (9%) expressed markers of profibrotic progenitor cells (CD9 high), which were observed previously and designated myofibrogenic progenitors in human and mice^19,20^ (**Figure 4C**). ASPC4 expressed PDE4D most highly.

**Figure 4.**
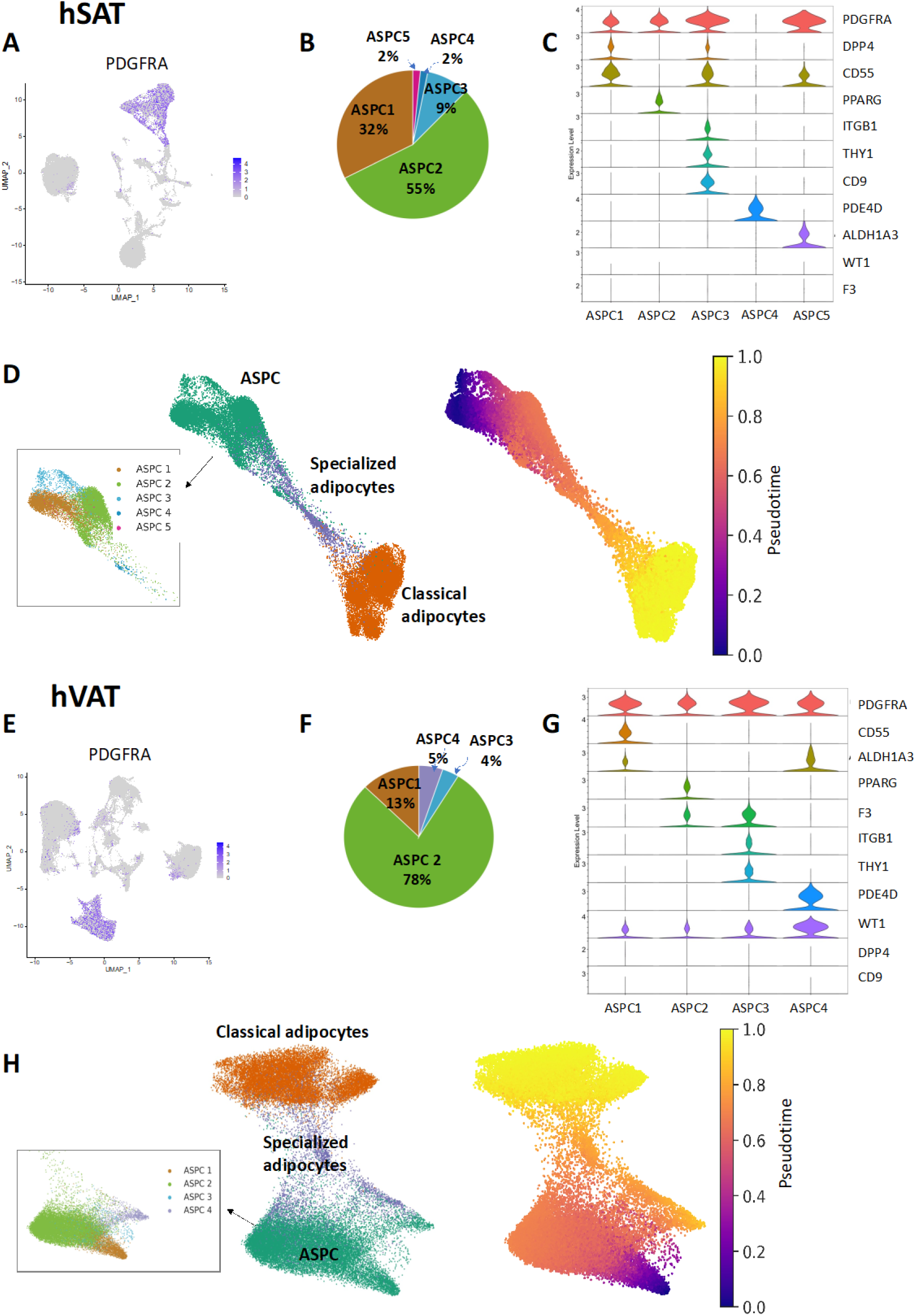
Differentiation paths from ASPCs to adipocytes in hSAT and hVAT. A. The expression of the ASPC-marker PDGFRA in UMAP representations of hSAT snNA-seq atlases. ASPCs constitute a well-defined cluster. B. The proportion of ASPC sub-clusters within the hSAT ASPCs. C. The expression of ASPC markers and of stem, fibrosis and adipogenic markers in hSAT ASPC sub-clusters. D. ASPCs, classical, and specialized adipocytes in hSAT colored by cell group **(left)** with inset showing ASPCs colored by sub-cluster. **Right**: Nuclei were colored by their differentiation pseudotime from ASPC stem cells (ASPC1). Specialized adipocytes were an intermediate state between ASPCs and classical adipocytes. Visualization was performed using ForceAtlas2 (FA) representation. E-F. Same as A-B for hVAT. G. The expression of ASPC markers and of stem, fibrosis, adipogenic and mesothelial markers in hVAT ASPC sub-clusters. H. Same as D for hVAT. Specialized adipocytes appear as an intermediate state between ASPCs and classical adipocytes. ASPC4 shows a pseudotime comparable to specialized adipocytes, suggesting it might lead to a different end-state.

Next, to follow the differentiation of ASPCs to adipocytes in hSAT we applied trajectory analysis^17^. This computational technique compares the RNA profiles of individual cells and orders them by the pseudotime of their differentiation from a given start state. As expected, a clear pseudotime path connecting hSAT-ASPC to adipocytes was observed (**Figure 4D**). In particular, the proliferative ASPC1, which was used as the start state for this analysis, and the profibrotic ASPC3 were the most distant from adipocytes, whereas ASPC2 and ASPC4 were the closest (**Figure 4D**). Importantly, the analysis uncovered that classical adipocytes are likely the terminal differentiation state of adipocytes, via specialized adipocytes (**Figure 4D**). The same path was observed with a different trajectory analysis tool^18^ (**Figure S3A**). This may suggest that the classical adipocyte differentiation state stems from loss, rather than gain, of specialized functions.

We repeated the above analysis for hVAT. *PDGFRA*-positive hVAT-ASPCs in this depot (**Figure 4E**) could be further segregated into four sub-clusters, denoted ASPC1-4 (**Figure 4F**). These clusters shared markers and integrated nicely with hSAT-ASPC1-4 (**Figure 4G**, **Figure S3B**). Trajectory analysis of hVAT ASPCs and adipocytes showed that, as in hSAT, the proliferative (ASPC1) and the profibrotic (ASPC3) clusters were the most distant from adipocytes, whereas the committed (ASPC2) was closer (**Figure 4H**). Furthermore, like in hSAT, hVAT-ASPCs linked to classical adipocytes via specialized adipocyte subtypes (**Figure 4H**), also upon using a different trajectory analysis tool (**Figure S3C)**. This is consistent with the notion that classical, rather than specialized adipocytes, are an ASPC differentiation end-state. Indeed, in both depots we observed a decrease in the expression of *PDGFRA* and two other ASPC-relevant genes from specialized to classical adipocytes, and an increase in the expression of adipocyte markers (**Figure S4**).

The above trajectory analyses also suggested that hVAT-ASPC might have an additional end-state for which ASPC4 is an intermediate step. Interestingly, ASPC4 relatively highly expressed mesothelial markers, such as WT1^21^ (**Figure 4H**). Hence, we extended the trajectory analysis to include hVAT mesothelial cells. Mesothelial cells could be divided into four sub-clusters, two of which were bona-fide mesothelial cells (Meso 1-2, that were mesothelin-high), and two that express the mesothelial markers MSLN and KRT19 at lower levels, but were nevertheless WT1-high (**Figure 5A**). The trajectory analysis showed that hVAT-ASPCs is linked to mesothelial cells via ASPC4 and through the sub-clusters that express mesothelial markers at lower levels, suggesting that these are ‘pre-mesothelial cells’ (pre-meso1,2, **Figure 5B**). Trajectories were supported also by alternative trajectory analysis^18^ (**Figure S5A**) and upon using negative controls **(Figure S5B**). Thus, at least two dominant differentiation paths could be envisioned for hVAT-ASPC, particularly the most distant subtype being hVAT-ASPC1: to adipocytes (specialized, then classical), and to mesothelial cells.

**Figure 5.**
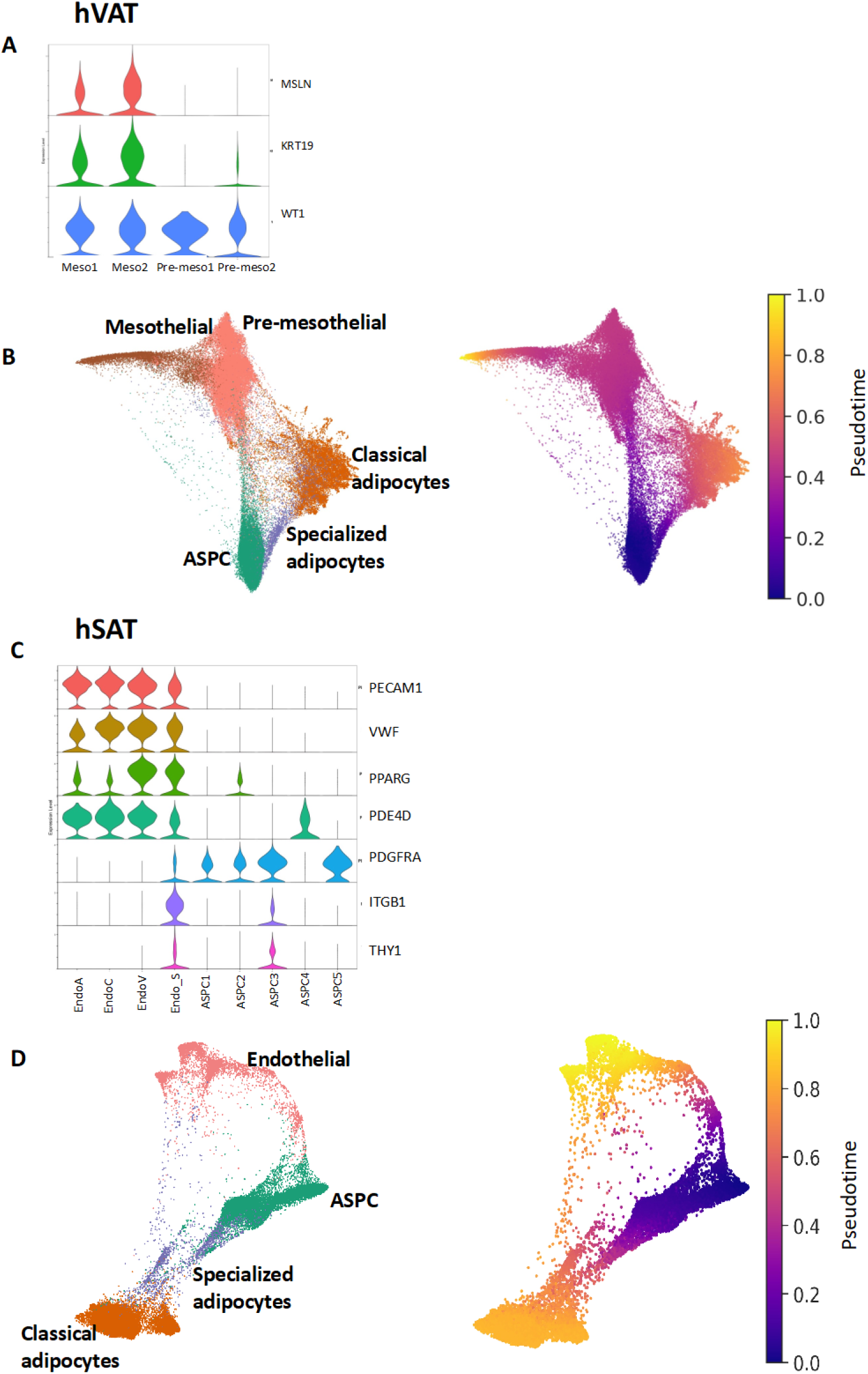
Potential depot-specific differentiation paths from ASPCs to adipocytes and other cell types. A. The expression of mesothelial markers in clusters of mesothelial and pre-mesothelial cells. B. ASPCs, pre-mesothelial, mesothelial, classical, and specialized adipocytes in hVAT colored by cell group **(left)**, and by their pseudotime from ASPC stem cells (ASPC1), **(right)**. Pseudotime analysis suggests that ASPCs could differentiate into classical adipocytes via specialized adipocytes, or to mesothelial cells via pre-mesothelial cells. C. The expression of endothelial and ASPC markers in clusters of endothelial and ASPCs. Endothelial and ASPCs clusters express distinct markers, apart from endothelial sub-cluster ‘Endo_S’ that expresses both types of markers. D. ASPCs, endothelial, classical and specialized adipocytes in hSAT colored by cell group **(left)**, and by their pseudotime from ASPC stem cells (ASPC1) **(right)**. Pseudotime analysis suggests two differentiation paths from ASPC, to either classical adipocytes via specialized adipocytes, or to of endothelial cells via Endo_S.

Given the bifurcated trajectory of hVAT-ASPC, we questioned whether hSAT-ASPCs could be similarly linked to an additional, dominant, differentiation end-state. We noticed that a cluster of endothelial cells, which was situated between other endothelial clusters and hSAT-ASPCs (**Figure 1B**), and similarly observed in ref.^10^ (cluster VC05), demonstrated expression of both endothelial (vWF) and ASPC (PDGFRA) markers (Endo_S, **Figure 5C**). This led us to test whether a transition state linking hSAT-ASPCs with endothelial cells is plausible. Trajectory analysis identified endothelial cells as another putative terminal differentiation state of hSAT-ASPC, particularly through the profibrotic ASPC3 (**Figure 5D**, **Figure S5C**). This was also supported by alternative trajectory analysis tool ^18^ and additional controls (**Figure S5C, S5D**, respectively).

Collectively, while the ASPC population exhibited at least two dominant trajectory pathways in both fat depots, the non-adipocyte terminal differentiation state differed, being endothelial in hSAT and mesothelial in hVAT.

### Adipocyte intercellular communication routes show fat-depot common and specific patterns

Adipose tissues are a main endocrine organ that heavily relies on paracrine communication by adipocytes. We started by considering the prototypical adipokines adiponectin and leptin, given that adipocytes are their only cellular source. Adiponectin and leptin showed a similar expression pattern among the various adipocyte subtypes in the two depots (except leptin was largely missing from VA6, **Figure S6A**). Adiponectin (ADIPOQ) is a ligand to 3 receptors that are transcribed from 3 distinct genes encoding *ADIPOR1, ADIPOR2* and *CDH13* (T-cadherin), while leptin signals through *LEPR*. We compared the expression of these genes among adipocyte subtypes in the two fat depots (**Figure S6A**). Among adiponectin receptors, *ADIPOR1* was largely missing in all adipocyte subtypes, while *ADIPOR2* seemed ubiquitous. Interestingly, angiogenic adipocytes in both depots (SA2, VA2) uniquely expressed the adiponectin receptor *CDH13* (**Figure S6A**). The leptin receptor *LEPR* was expressed in most subtypes. Expanding the view to the expression of adiponectin and leptin receptors in other cell types, revealed that expression of *ADIPOR2* is detectable in more cell types in hVAT than in hSAT (**Figure S6B**), including in cell types that are present in both depots (see, for example, mast cells and endothelial cells). Interestingly, *CDH13* expression characterized endothelial cells in both depots (though lymphatic endothelial cells expressed *CDH13* only in hSAT), suggesting that adiponectin may similarly affect angiogenic adipocytes and endothelial cells.

Next, we considered whether adipocyte subtypes show distinct and/or depot-differential communication patterns between them. We first estimated the total number of intercellular interactions among adipocyte subtypes using CellPhoneDB (**Table S8**). Inter-adipocyte subtypes communication seemed more active in hSAT than in hVAT (median number of interactions 212 versus 110, respectively, **Figure 6A**). Next, we estimated communication probabilities for specific intercellular ligand-receptor pairs using CellChat, which also identifies “key” incoming and outgoing signaling pathways^22^. This analysis suggested that classical adipocytes (SA1, VA1) were the main senders of adiponectin in both depots, and of leptin in hVAT (**Figure 6B**), possibly reflecting their dominant proportions among the various adipocyte subtypes. While adiponectin signaling was a key pathway in both depots (**Figure 6B**), leptin-leptin receptor communication was a key pathway only in hVAT (**Figure 6C**), consistent with low *LEPR* expression in SA1 (**Figure S6A, Table S9**). This suggests that while hSAT is the major source of leptin production ^23,24^, it might be less sensitive to leptin.

**Figure 6.**
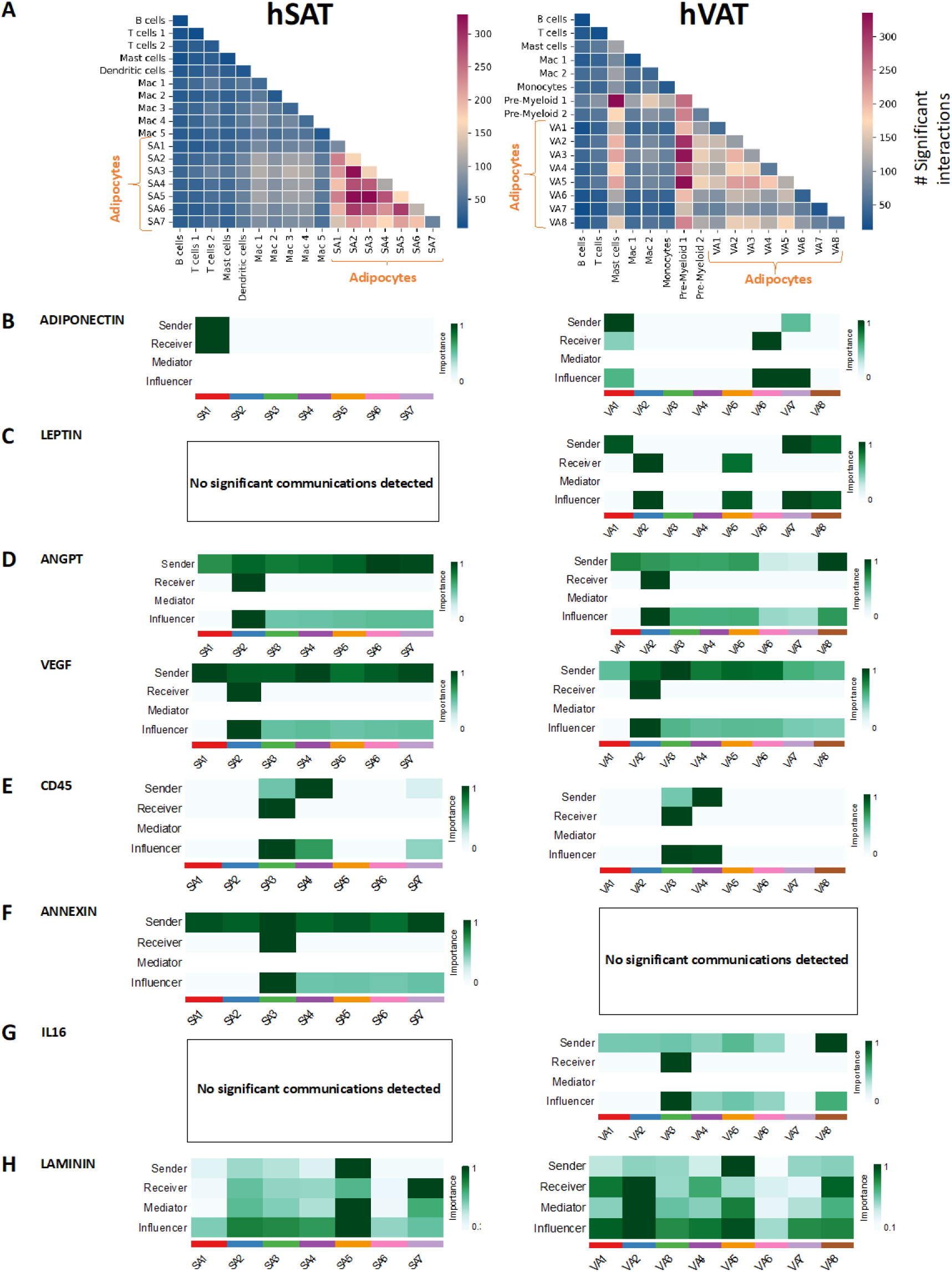
Adipocytes and immune cells in SAT and VAT intercellular and signalilng communication patterns. A. The total number of interactions between cell types in SAT and VAT. The number of interactions among adipocyte subtypes is higher in hSAT. The number of interactions between adipocytes and the macrophage/myeloid compartment or mast cells is higher in hVAT compared to hSAT. Heatmap was calculated using CellPhoneDB. B-I. The senders and receivers of specific signaling pathways among adipocytes in hSAT and hVAT. Analysis was performed using CellChat. B. Classical adipocytes are the main senders on Adiponectin in both depots. C. Leptin pathway is missing from hSAT. In hVAT, classical adipocytes VA1 and VA7,VA8 are the main senders and VA2 and VA5 are main receivers. D. In both depots, angiogenic adipocytes SA2,VA2 are the main receivers of the angiogenesis-related ANGPT and VEGF signaling pathways. E. In both depots, immune-related adipocytes SA4,VA4, and SA3,VA3, are the main senders and receivers, respectively, of the immune CD45 signaling pathway. F. In hSAT alone, the anti-inflammatory signaling pathway ANNEXIN is received mainly by SA3. G. In hVAT alone, the pro-inflammatory signaling pathway IL16 is received mainly by VA3. H. In both depots, the ECM adipocytes SA5,VA5 are the main senders of the ECM-related laminin signaling pathways.

Analysis of additional signaling pathways revealed several adipocyte subtype-specific patterns. In both depots, angiogenic adipocytes (SA2, VA2) were the most likely adipocyte subtype to be targeted by the angiogenesis-related ANGPT and VEGF signaling pathways (**Figure 6D)**. Immune-related adipocyte subtypes (SA3,4 and VA3,4) similarly exhibited CSF and CD45 immune pathways as key communication patterns (**Figure 6E, Table S9**). However, while the hSAT SA3 adipocytes were targeted by the anti-inflammatory annexin key pathway (**Figure 6F)** and galectin (**Table S9**), their respective hVAT equivalents VA3 were targeted by the pro-inflammatory IL16 key pathway (**Figure 6G**). This is consistent with the notion of VAT being more inflammatory than SAT, suggesting that these adipocyte subtypes participate in mediating this depot difference. ECM adipocytes (SA5, VA5) were the main senders of the laminin pathway, which plays major roles in adipose tissue ECM deposition, in both depots (**Figure 6H**).

We further explored whether the hSAT-hVAT difference in inflammatory communication extends beyond inter-adipocyte interactions, by considering communication with the adipose macrophage/myeloid compartment (including bona-fide macrophages, dendritic cells, monocytes, and “pre-myeloid cells”). In hSAT, the numbers of adipocyte subtype interactions with this cell compartment were significantly lower than inter-adipocyte interactions (Mann-Whitney test p<E-9, **Figure 6A**). This was in contrast to hVAT, in which adipocytes interaction with subclasses of this compartment (pre-myeloid 1,2) seemed to be more intense than the inter-adipocyte interactions (Mann-Whitney test p=0.002, **Figure 6A**). Similarly, mast cells interacted more with adipocytes and with myeloid cells in hVAT than in hSAT (**Figure 6A**). To further explore this seeming more pro-inflammatory pattern in hVAT compared to hSAT, we examined the annexin and IL16 pathways upon including the macrophage/myeloid compartment. The anti-inflammatory annexin signaling was much more pronounced in hSAT versus hVAT, and involved many more cell types as senders or receivers (**Figure S7, Table S9**). The pro-inflammatory IL16 signaling involved many cell types of the macrophage/myeloid compartment in both depots and with similar intensity, except for the unique role of VA3 as the sole adipocyte subtype receiver.

Another difference between the depots’ communication patterns was the low interaction intensity between adipocytes and mast cells in hSAT compared to hVAT (**Figure 6A**). Mast cells were targeted by adiponectin in hVAT but not in hSAT (**Figure S6B**). Likewise as source, they were highly involved in ECM, fibrosis and IL16 pathways in hVAT (**Figure S8**). Thus, depot-differential communication patterns between adipocyte subtypes and mast cells also contribute to the differential inflammatory and fibrotic “tone” between the two fat depots.

## DISCUSSION

Single-nucleus RNA sequencing has recently been employed by several groups to describe the cellular landscape of human adipose tissues with unprecedented resolution^9–13,15^. These adipose tissue cellular “atlases” enabled to unbiasedly identify previously unrecognized diversity and subtypes of each of the different cell types comprising adipose tissue, estimate inter-cell-type communication patterns, and increase our understanding of how adipose tissue cell composition may vary in the population. Interestingly, when comparing hSAT to hVAT, although some fat depot differences were noted^9,10,12,15^, overall the cellular landscape based on the nuclear transcriptome profiling largely revealed similarities between the depots. This is despite large body of evidence demonstrating that hSAT and hVAT are functionally, and even transcriptionally (by whole-tissue, bulk RNA sequencing), distinct.

In the present study, we hypothesized that in-depth depot-specific analysis of adipocyte atlases obtained from sNucSeq profiling could shed light on some of the differences between human fat depots. Our main findings relate to adipocyte subtype functionality, differentiation trajectories from ASPCs, and/or inter-cellular communication patterns, as detailed below.

Both hSAT and hVAT harbor a majority of “classical adipocytes”, and several “specialized adipocyte” subtypes, each enriched in unique functions (**Figure 3**). Consistent with previous studies, the adipocyte subtype composition is indeed relatively similar between hSAT and hVAT (**Figure 2A**). The in-silico characterization of the unbiasedly-identified adipocyte subpopulations relied on pathway enrichment analyses, which, relative to individual gene markers, provided a clearer and more robust biologically-relevant annotation (**Figure 3A-F**). In both depots classical adipocytes constituted the majority of adipocyte subtypes, while the next-sized adipocyte subgroups (with depot-different abundances) were ECM-related adipocytes, angiogenic adipocytes, and immune-regulating adipocytes (**Figure 2A**). These adipocyte subpopulations were identifiable among hSAT and hVAT adipocytes in an independent published dataset (**Figure 3G-L**). Previously, three hSAT adipocyte subtypes were identified by different expression level of *LEP, PLIN1* and *SAA,* one of which responded transcriptionally to in-vivo insulin stimulation^11^. While our data cannot test for insulin sensitivity, marker genes used in this study suggest that their parallel adipocyte subtypes likely fall in our data within the large, classical adipocyte cluster. This cluster, by our analysis, only mildly (compared to specialized adipocytes) preferentially expressed lipid metabolism and handling genes. Furthermore, all adipocyte subtypes in our study similarly expressed *LEP, ADIPOQ, PLIN1* and other adipocyte-unique genes, attesting to their adipocytic character. Depot-specific adipocyte subtypes were identified in ref. ^9^, and consistently, a small hVAT-unique adipocyte subtype, VA6, was observed in our dataset as well. This adipocyte cluster was characterized by relatively high expression of ribosomal genes, possibly consistent with a high protein translation activity that characterizes hVAT (**Figure 3F**).

Trajectory analysis from ASPCs to adipocytes suggested that in both depots classical adipocytes are the terminal differentiation state, via specialized adipocytes (**Figure 4**). This suggests that unique functions of specialized adipocytes are “lost” during biogenesis of classical adipocytes. hSAT and hVAT adipocytes likely emerge from an ASPC sub-population that expresses PPARG. In contrast to this between-depot similarity, the more proliferative and likely more pluripotent ASPC subtype can give rise to endothelial cells in hSAT, and to mesothelial cells in hVAT (**Figure 5**). Our insights into ASPC to adipocyte differentiation trajectories are consistent with several trends observed in previous studies, that suggest divergent trajectory paths from ASPCs (^10,12^). Considering the ASPC (FAP) population, Massier et al., ^10^ demonstrated in hSAT a bifurcated trajectory arising from ASPC subpopulations with different commitment towards adipogenic differentiation. Considering ASPCs from white and brown human fat, Palani et al., ^1^ also identified a bifurcation of differentiation trajectories into adipocytes and Wnt-regulated adipose tissue-resident (SWAT) cells. We included into the trajectory analysis also terminally differentiation states, with adipocytes being an obvious trajectory target of ASPCs committed to adipogenesis, which we identified in both depots. Depots differed in the putative second dominant terminal differentiation state, with endothelial cells in hSAT and mesothelial cells in hVAT. Historically, adipocytes were proposed to differentiate from mesothelial cell origin^25^, however, this pathway was more recently deemed less likely in mice^21^. Mesothelial cells are a cell population unique to VAT^9,10^. However, we noted that the bona-fide *MSLN*-positive cells were close to a large cell population that was *MSLN-*negative, but which nevertheless expressed other mesothelial, but not ASPC markers (i.e., “pre-mesothelial cells”). Indeed, the ASPC-to-mesothelial cell trajectory we observed in hVAT passed via this cell population, consistent with their intermediate state. In hSAT, the non-adipogenic ASPC trajectory arm leading to endothelial cells passed through a subpopulation of cells co-expressing endothelial and ASPC markers. This subpopulation was also observed in Massier et al.^10^. Jointly, our analysis further elucidates putative depot-specific differences in differentiation trajectories of the human adipose stem cell compartment.

As for intercellular communication patterns, hSAT adipocyte subtypes were predicted to highly interact, more than with adipose tissue immune (mostly myeloid) cells, while this is opposite in hVAT (**Figure 6A**). Moreover, adipocyte sub-populations common to the two depots, such as immune-regulating adipocytes (SA3, VA3), participate in inter-cellular communication through depot-different dominant interaction pathways (**Figure 6F-G**). These provide new ways to understand the more pro-inflammatory nature of hVAT over hSAT.

Our study has several noteworthy weaknesses and strengths. Common to our and other single-cell studies are the limited number of samples assessed (mainly a result of the high cost of the technology), and indeed, “meta-analysis” of data from different centers has recently been published^10^. We employed several approaches to address this shortcoming: First, donors present diversity of sex, age, and BMI(**Figure 1A**), while carefully selected to exclude severe morbid conditions. We integrated our adipocyte dataset with that of Emont et al., to verify that adipocyte subtypes in our data are also identifiable in an independent dataset (**Figure 3G-L**). Finally, we utilized an in-house deconvolution algorithm tailored for snRNA-seq of human adipose tissue (sNuConv ^16^), to validate key findings, including the identification of classical and specialized adipocytes in a larger cohort (**Figure S2**). Another important aspect of our study is the separate analysis of each depot, and particularly of the adipocyte population. While combined analysis of data from both depots could enhance signals that might otherwise remain undetected, it might also mask some of the differences. The independent analysis of each depot allowed us to find commonalities and differences in an unbiased manner. These were then verified using alternative computational analyses, including machine learning classification and deconvolution, and additional datasets, including other snRNA-seq and bulk RNA-sequencing profiles of human adipose tissue.

In conclusion, by separately analyzing the cellular landscape of hSAT and hVAT, we refine understanding of the commonalities and differences between human adipose tissue depots at the level of adipocyte cell composition, differentiation trajectory pathways, and inter-cellular communication.

## METHODS

### Collection of human AT samples

AT biopsies with at least 1gr were obtained from adult patients without severe morbid states or obesity complications who had signed in advance a written informed consent before undergoing elective abdominal surgeries. Five patients donated SAT and VAT samples (paired). Tissues were immediately delivered from the operating room to the lab, where they were snap frozen in liquid nitrogen and stored at −80°C, as detailed previously^26,27^. For 14 samples, ethical approval of the study procedures was obtained by the Helsinki Ethics Committee of Soroka University Medical Center (approval no: 15-0348). For one sample, collection was approved in the context of the Leipzig Obesity BioBank (LOBB; n>8,000 donors; BMI range 13-129kg/m^2^; age range: 18-96 years) by the Ethics Committee of the Medical Faculty of the University of Leipzig (approval numbers: 159-12-21052012 and 017-12-ek) and performed in accordance with the Declaration of Helsinki. Clinical characteristics of donors and samples are presented in **Figure 1A** and **Table S1**.

### Single-nucleus RNA Sequencing

Human AT samples were analyzed by isolating nuclei, barcoding their transcriptome using Chromium 10X technology, generating cDNA and then expression libraries followed by sequencing (of the expression libraries). Briefly, nuclei were isolated from 500 mg of frozen hSAT or hVAT samples with all steps carried out on ice. One ml of ice-cold adipose tissue nuclei lysis buffer (AST: 5 mM PIPES, 80 mM KCl, 10 mM NaCl, 3 mM MgCl2 and 0.1% IGEPAL CA-630) supplemented with 0.2U/µl of RNAse Protector (Roche, Switzerland), was added to each sample, followed by mincing the frozen tissue using small surgical scissors. Then, small pieces of minced tissue in 1 ml of AST lysis buffer were transferred into 7 ml WHEATON® Dounce Tissue Grinder (DWK Life Sciences, Germany) and additional 2 ml of AST were added. Tissues were dissociated to retrieve nuclei by 12 strokes with loose pestle (A), followed by 10 strokes with tight pestle (B). Samples were then incubated for 7 minutes on ice to ensure maximal nuclei retrieval, filtered through 70 µm SMARTStrainer (Miltenyi Biotech, Germany), and centrifuged (4°C, 5 minutes, 500 RCF). Supernatants were discarded, and nuclei pellet was resuspended in 2.5 ml of ice-cold PBS 0.5% BSA, supplemented with 0.2U/µl of RNAse Protector, and filtered through 40 µm SmartStrainer, followed by additional centrifugation. Next, 2 ml of supernatant were discarded, and nuclei pellet was resuspended in the remained 0.5 ml of ice-cold PBS 0.5% BSA. Nuclei were counted using LUNA-FL™ Dual Fluorescence Cell Counter (logos biosystems, South Korea), and 13,000-16,500 nuclei/sample were immediately loaded onto 10X Chromium Controller (10X GENOMICS, CA, USA) according to the manufacturer’s protocol (Chromium Next GEM Chip C. Chromium Next GEM Single Cell 3’ Kits v3.1). cDNA and gene expression libraries size fragments were assessed using 4150 TapeStation System (Agilent, CA, USA) with high sensitivity D5000 ScreenTape, while their concentrations were quantified using Qubit 4 Fluorometer using Qubit dsDNA HS Assay Kit (ThermoFisher Scientific, MA, USA). Gene expression libraries were sequenced by NextSeq 550 using High Output Kit v2.5 (150 Cycles) kit and NovaSeq 6000 using S1-100 cycles flow cell kit at sequencing depth of at least 50,000 reads per nucleus.

### snRNA-seq data pre-processing and quality control

Per sample, reads were transformed into a raw counts matrix with 10x Genomics CellRanger software^1^ 6.0.1, which was used as input for CellBender^28^ to remove ambient RNA. For downstream analysis, only cell barcodes that were determined to be true cells by both CellRanger and CellBender algorithms were used. The resulting raw counts matrix of each sample was analyzed with Seurat v3.0 R package^29,30^. Our quality control procedure included removal of broken nuclei, i.e., nuclei with less than 200 genes or nuclei that contained more than 20% mitochondrial genes. Counts were normalized and scaled using the 3,000 most variable genes. Linear dimensionality reduction via principal component analysis (PCA) and graph-based clustering via k-nearest-neighbors (KNN, k=20) methods were used to group individual cells into cell subsets. In order to remove plausible doublets, we removed cell clusters with low mean gene count and low mean reads counts (less than one standard deviation below the sample’s mean), and then applied DoubletFinder R package^31^. All cells that were marked as doublets by DoubletFinder were removed, along with cell subsets that contained more than 65% doublets. Once each sample passed quality control, we integrated all samples using Harmony^32^ and performed dimensionality reduction and clustering again. Clusters were calculated with 0.8 resolution. To avoid over-clustering, clusters were tested with AssessNodes function from Seurat v2.0, which we restored to fit Seurat V3.0 objects. Clusters that were not significantly different were merged (p<0.05). To avoid sample-specific clusters, clusters derived mostly from a single sample (over 80% of cluster cells) were not considered in subsequent analyses.

### Identification of cell clusters and cluster annotation

Differentially expressed marker genes were calculated using Seurat v3.0 FindAllMarkers function (logfc.threshold=-inf and min.pct=0). Cell subsets were annotated using known markers from the literature, expert knowledge, and CellTypist R package^33^. Detailed description of the annotation of non-adipocytes appears in ref.^16^.

### Adipocyte subtype detection and characterization

Adipocyte clusters were extracted from the atlas of each depot and re-analyzed in a depot-specific manner, including normalization, determination of variable genes, scaling, and integration using Harmony^32^. To characterize adipocyte subtypes we used gene set enrichment analyses (GSEA) with clusterProfiler^34^ with GO as reference gene sets. We then associated an adipocyte subtype with a subgroup of processes for which it was significantly enriched (adjusted p<0.05, selected processes appear in **Table S6**). For visualization of the enrichment for a given process subgroup, we calculated per subtype the sum of the GSEA normalized enrichment score (NES) for this subgroup (if no enrichment was detected the NES value was set to zero). Cells within a subtype were then colored by the calculated score of the subtype.

### Comparison between SAT and VAT adipocytes

Adipocytes were extracted from SAT and VAT. The combined dataset was re-analyzed, including normalization, determination of variable genes, scaling, and integrated using Harmony^32^ with the grouping variable set to sample name and depot.

### Comparison between in-house and other SAT and VAT adipocyte datasets

Adipocytes were extracted from Emont et al.^9^ and were combined with adipocytes detected in this study per depot. The combined dataset was re-analyzed, including normalization, determination of variable genes, scaling, and integration using Harmony^32^ with the grouping variable set to tissue source (in-house or Emont et al.) and sample name. Next, to avoid over-clustering, clusters were tested using AssessNodes function from Seurat v2.0 (p-values 0.08 for hSAT and 0.09 for hVAT). Integrated clusters were annotated based on the dominant adipocyte subtype contributing the majority of its cells to the cluster. We also used machine-learning to test for the presence of adipocyte subtypes in the Emont et al dataset^9^. Per depot, we trained a separate random forest classifier per in-house adipocyte subtype with over 200 nuclei. Training was based on the top 20 positive differentially expressed genes for each adipocyte subtype that were also included in Emont et al. dataset^9^. Training was assessed using 5-fold cross validation on 80% percent of the data, and then tested on the remaining data. The auROC and auPRC of each model appear in **Figure S2A**. Next, we applied each trained model to classify adipocytes of the corresponding depot from ref.^9^. Adipocytes were annotated to the subtype with the highest probability score above 0.2. The prediction score of each model per depot appears in **Table S7**.

### Deconvolution analysis

Analysis was performed using sNuConv tool^16^. We trained a separate model for SAT and VAT, based on five SAT and seven VAT samples for which bulk RNA-sequencing data were available^16^. Per depot we trained two separate models, one model treating adipocytes as a single cell-type, and the other model distinguishing between classical and specialized adipocytes. The performance of trained models was assessed using leave-1-out, by comparing the estimated proportions and true, snRNA-seq-derived proportions of the different cell types in the left-out sample. Pearson r coefficient was used to quantify the correlation accuracy per model. Trained models obtained r≥0.88 in SAT and r≥0.93 in VAT. Trained models were applied using the sNuConv tool to 73 paired bulk RNA-seq profiles of human adipose tissue. Paired omental visceral and abdominal subcutaneous samples were obtained from 73 donors that have been selected from the Leipzig Obesity BioBank (LOBB). Patients included into the deconvolution analysis had a preoperative BMI of 54.5 ± 9.3 kg/m2 and age of 44.1 ± 9.2 years. Type 2 diabetes (T2D) was diagnosed in 28 patients. For each individual, adipose tissue was collected during elective laparoscopic abdominal surgery and anthropometric as well as laboratory measurements were obtained, as previously described^35^.

### Trajectory analysis

Trajectory analysis was conducted using Palantir^17^ version 1.2.0 with the parameters set as in the Palantir tutorial, and using DPT^18^ as implemented in Scanpy^36^ version 1.9.3 (dpt function). Per analysis, the nuclei corresponding to the relevant cell types were extracted and reanalyzed, including normalization, log-transformation, detection of highly variable genes, dimensionality reduction, and integration using Harmony. Analyses were based on the 1,500 most variable genes, as suggested by Palantir, except for the analysis of VA-ASPC, where the number of most variable genes was raised to 2,500, in order to obtain output. The visualization of trajectories was done using ForceAtlas2^37^. The start cell for each analysis in both SAT and VAT was an ASPC1 nucleus with zero F3 expression that was randomly selected from the 100 cells with highest DPP4 expression. For visualization of gene expression patterns along the trajectory we used MAGIC imputation with default parameters^38^.

### Intercellular communication analysis

Prediction of enriched intercellular interactions were performed using CellphoneDB^39^ version 4.0.0 with default parameters. The CellphoneDB heatmap was obtained using ktplotspy library. For visualization, we summed the ligand-receptor interactions of each pair of cell-types (i.e VA2->VA4 and VA4->VA2) and plotted it using Seaborn library. The datasets created by CellphoneDB appear in **Table S8**. Prediction of intercellular communication probabilities and key signaling pathways were performed using CellChat version 1.6.0^40^ with default parameters, except population.size was set to True to minimize the effects of larger cell groups in the data. Analyses were performed per depot, using either adipocytes alone or adipocytes and adipose immune cells. Key signaling pathways were detected using netAnalysis_signalingRole_network function. Visual representation of inferred communication network was obtained using netVisual_aggregate function with chord layout and netVisual_heatmap for heatmap representation. Significant communication interactions and key signaling pathways appear in **Table S9.**

## ACKNOWLEDGEMENT

This publication is part of the Human Cell Atlas – www.humancellatlas.org/publications/. This study has been made possible in part by CZI grant CZIF2019-002441 and grant DOI https://doi.org/10.37921/206883bpivjy from the Chan Zuckerberg Initiative Foundation (funder DOI 10.13039/100014989).

## SUPPLEMENTARY FIGURES

**Figure S1.**
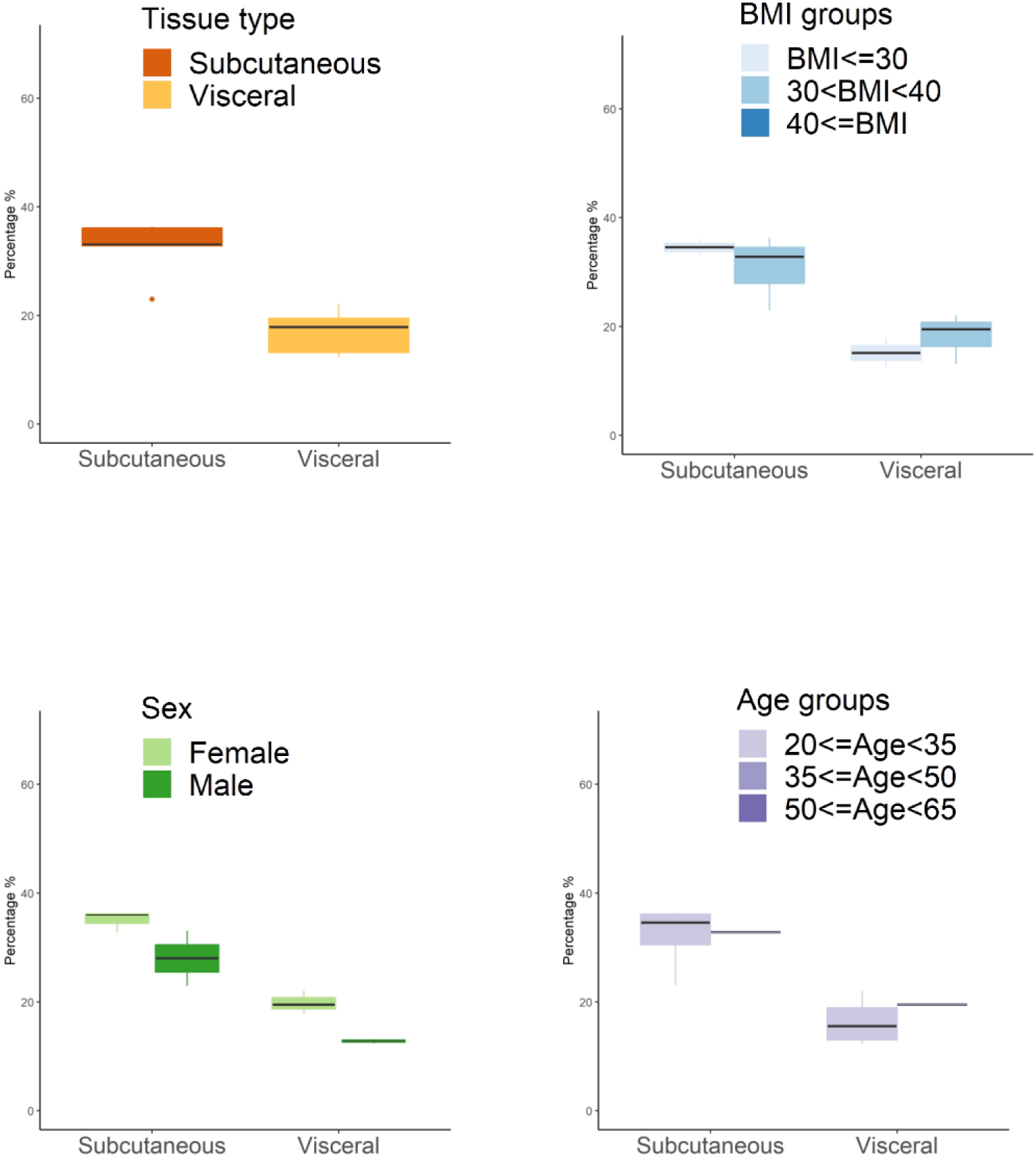
The fraction of adipocytes per depot, stratified by sex, age, and BMI in the five paired samples.

**Figure S2.**
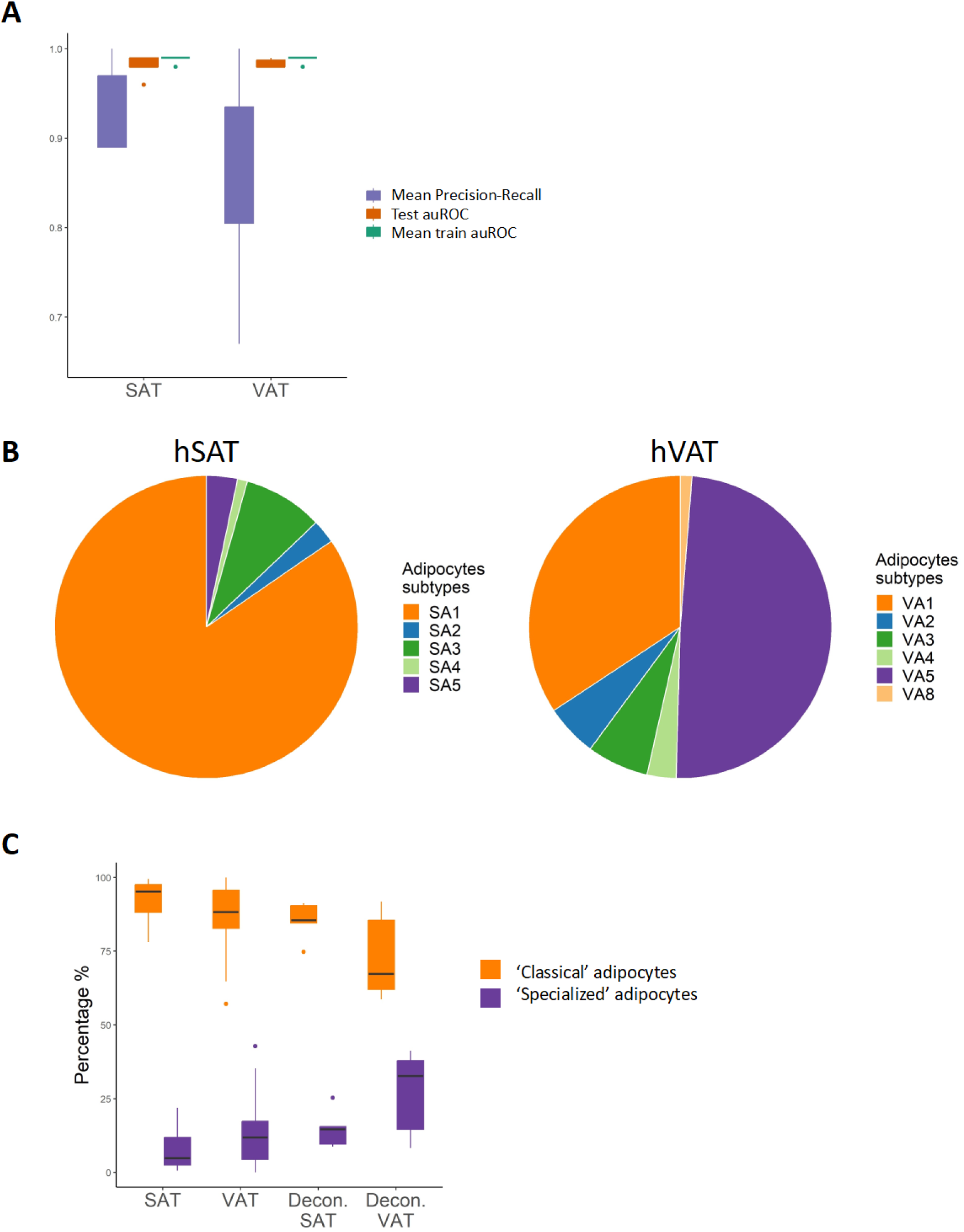
Identification of classical and specialized adipocytes in additional datasets. A. The performance of the machine-learning classifiers of adipocyte subtypes. Shown are the area under the receiver operating characteristic curve (auROC) in the five-fold cross validation training (mean train auROC) and in the testing (test auROC), and the mean area under the precision recall curve (mean auPRC) for the different adipocyte subtype classifiers. B. The proportions of adipocyte subtypes in the Emont et al. dataset^9^ predicted using classifiers. C. Proportions of classical and specialized adipocytes, calculated from the total number of adipocytes per depot being 100%. Shown are the ‘true’ proportions in our in-house snRNA-seq data (SAT, VAT) and estimated proportions using deconvolution analysis of 73 paired bulk RNA-seq profiles (‘Decon.’ SAT and VAT).

**Figure S3.**
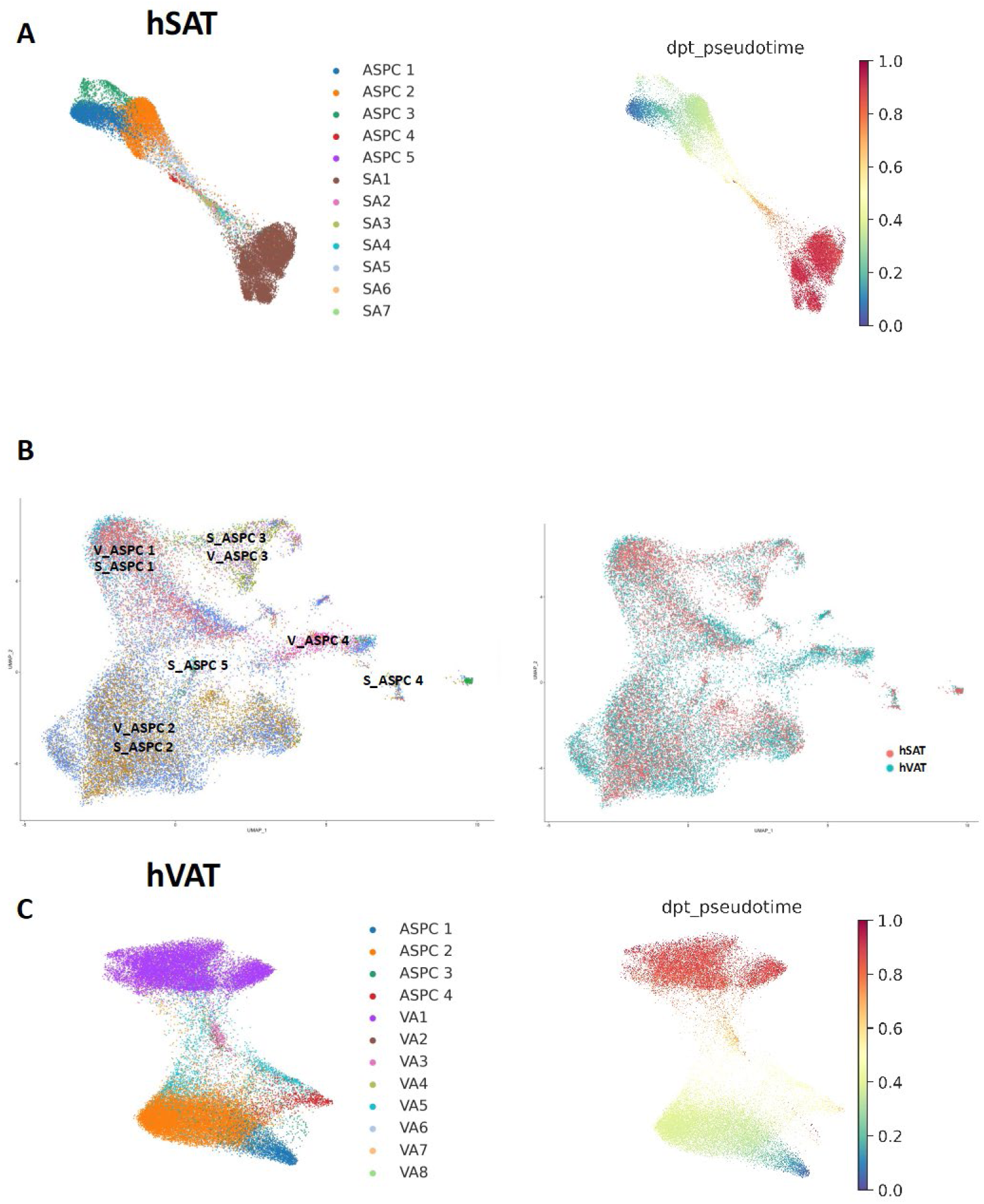
Differentiation paths from ASPCs to adipocytes in hSAT and hVAT. A. ASPC and adipocyte sub-types in hSAT colored by cell group (**left**), and by their differentiation pseudotime from ASPC stem cells (ASPC1) according to DPT (**right**). Specialized adipocytes were an intermediate state between ASPCs and classical adipocytes. B. Integration of ASPCs from hSAT and hVAT, colored by sub-cluster (**left**) and by dept (**right**). The respective ASPC1-4 clusters from hSAT and hVAT clustered together. C. Same as A for hVAT. Again, specialized adipocytes were an intermediate state between ASPCs and classical adipocytes. Nuclei maps were represented using ForceAtlas2 (FA).

**Figure S4.**
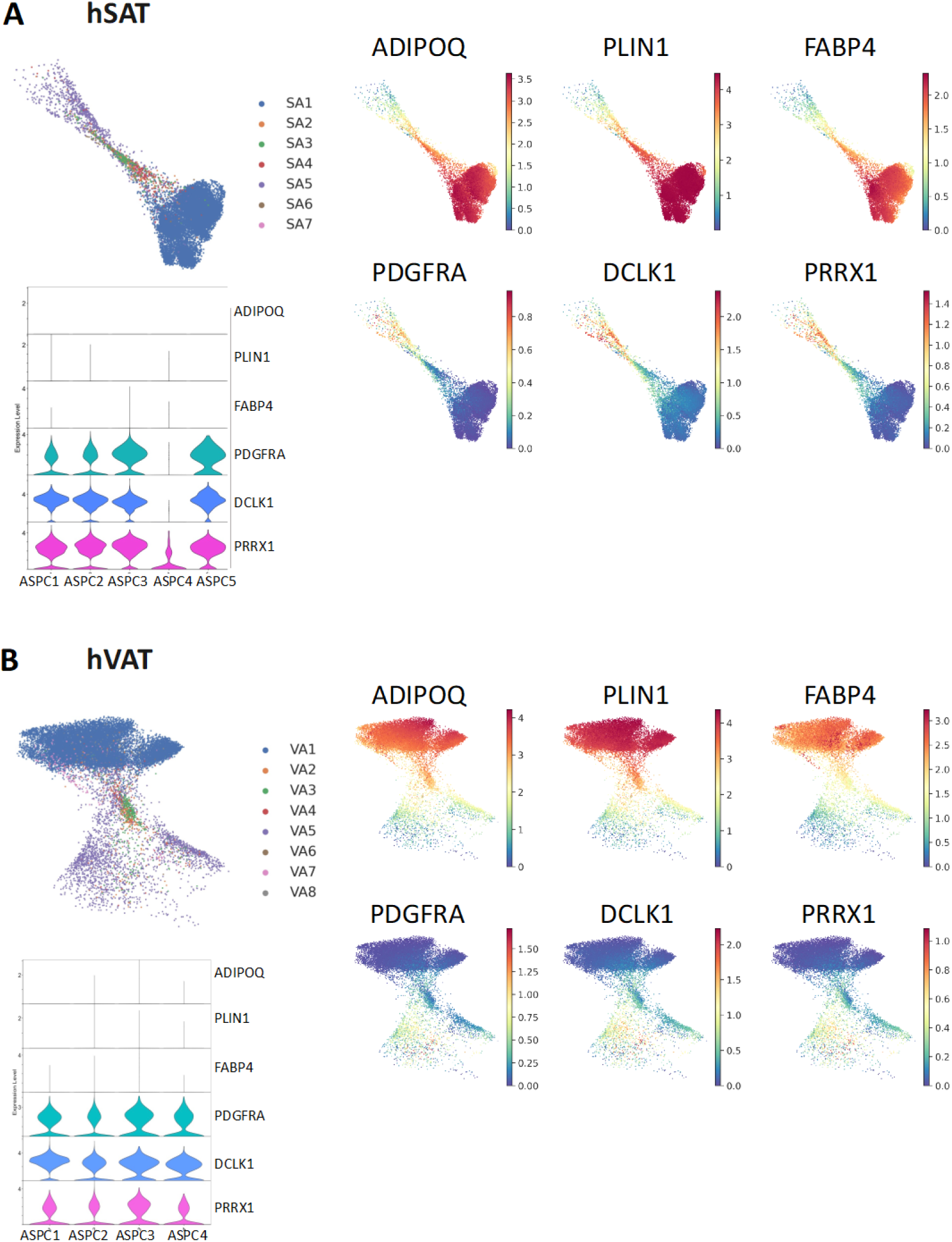
The expression ASPC and adipocyte relevant genes in the different adipocyte subtypes. In hSAT (A) and hVAT (B), the top and bottom rows show adipocyte and ASPC relevant genes, respectively. The expression of adipocyte relevant genes increases from specialized to classical adipocytes, whereas the expression of ASPC relevant genes decreases. Violin plots show the expression of these genes in ASPCs.

**Figure S5.**
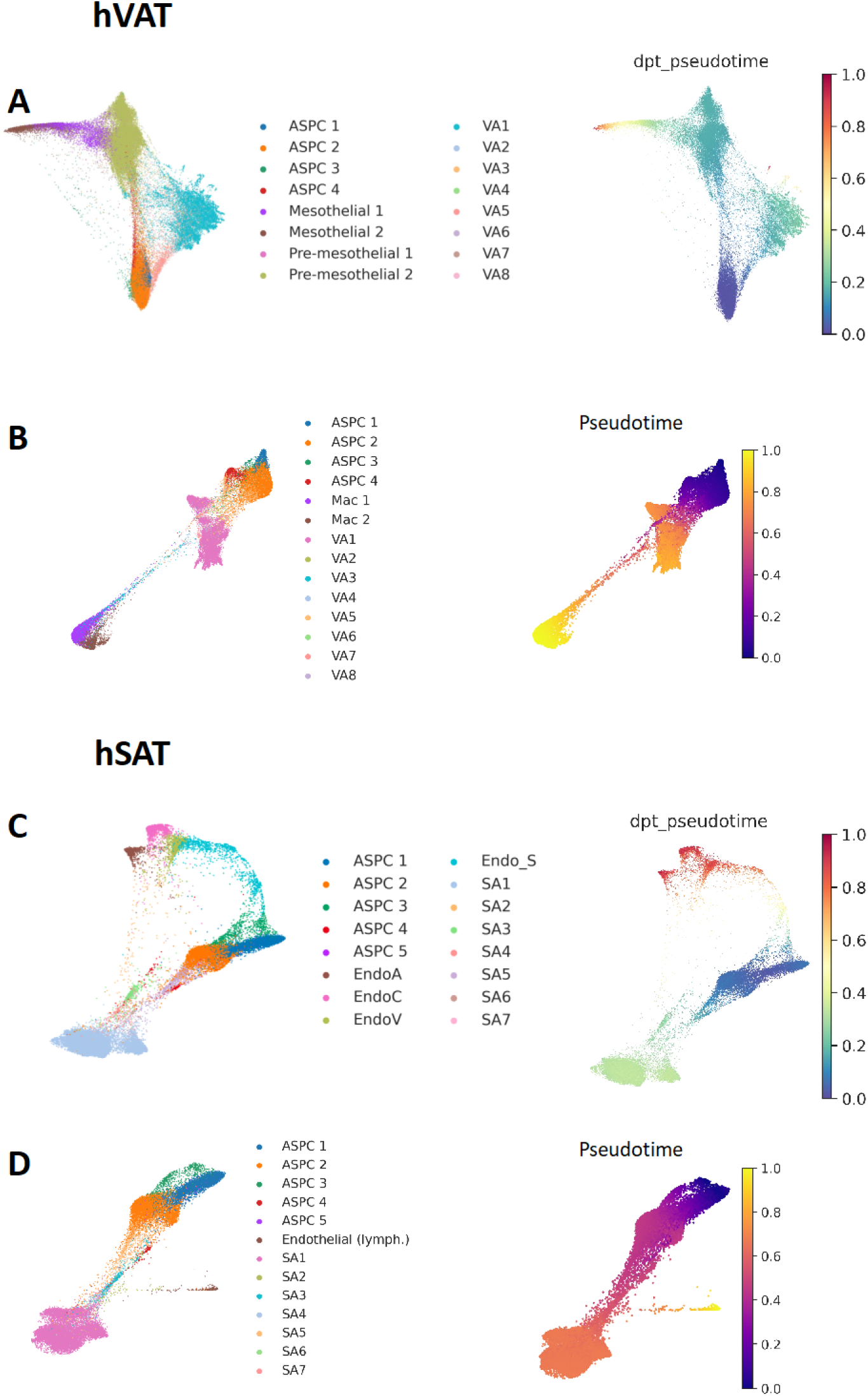
Potential depot-specific differentiation paths from ASPCs to adipocytes and other cell types. A. hVAT ASPC, adipocyte, and mesothelial sub-types colored by cell group (**left**), and by their differentiation pseudotime from ASPC stem cells (ASPC1) according to DPT (**right**). This alternative analysis suggests that ASPCs could differentiate into classical adipocytes via specialized adipocytes, or to mesothelial cells via pre-mesothelial cells, consistent with Figure 5B. B. hVAT ASPC, adipocyte, and macrophage sub-types colored by cell group (**left**), and by their differentiation pseudotime from ASPC stem cells (ASPC1) according to Palantir (**right**). The trajectory illustrates a long and scattered differentiation path from ASPCs to macrophages via adipocytes, hence acting as a negative control for Figure 5B. C. hSAT ASPC, adipocyte, and endothelial sub-types colored by cell group (**left**), and by their differentiation pseudotime from ASPC stem cells (ASPC1) according to DPT (**right**). This alternative analysis suggests two differentiation paths from ASPC, to either classical adipocytes via specialized adipocytes, or to of endothelial cells via Endo_S, consistent with Figure 5D. D. hVAT ASPC, adipocyte, and lymphatic endothelial sub-types colored by cell group (**left**), and by their differentiation pseudotime from ASPC stem cells (ASPC1) according to Palantir (**right**). The trajectory illustrates a scattered differentiation path from ASPCs to lymphatic endothelial via specialized adipocytes, hence acting as a negative control for Figure 5D.

**Figure S6.**
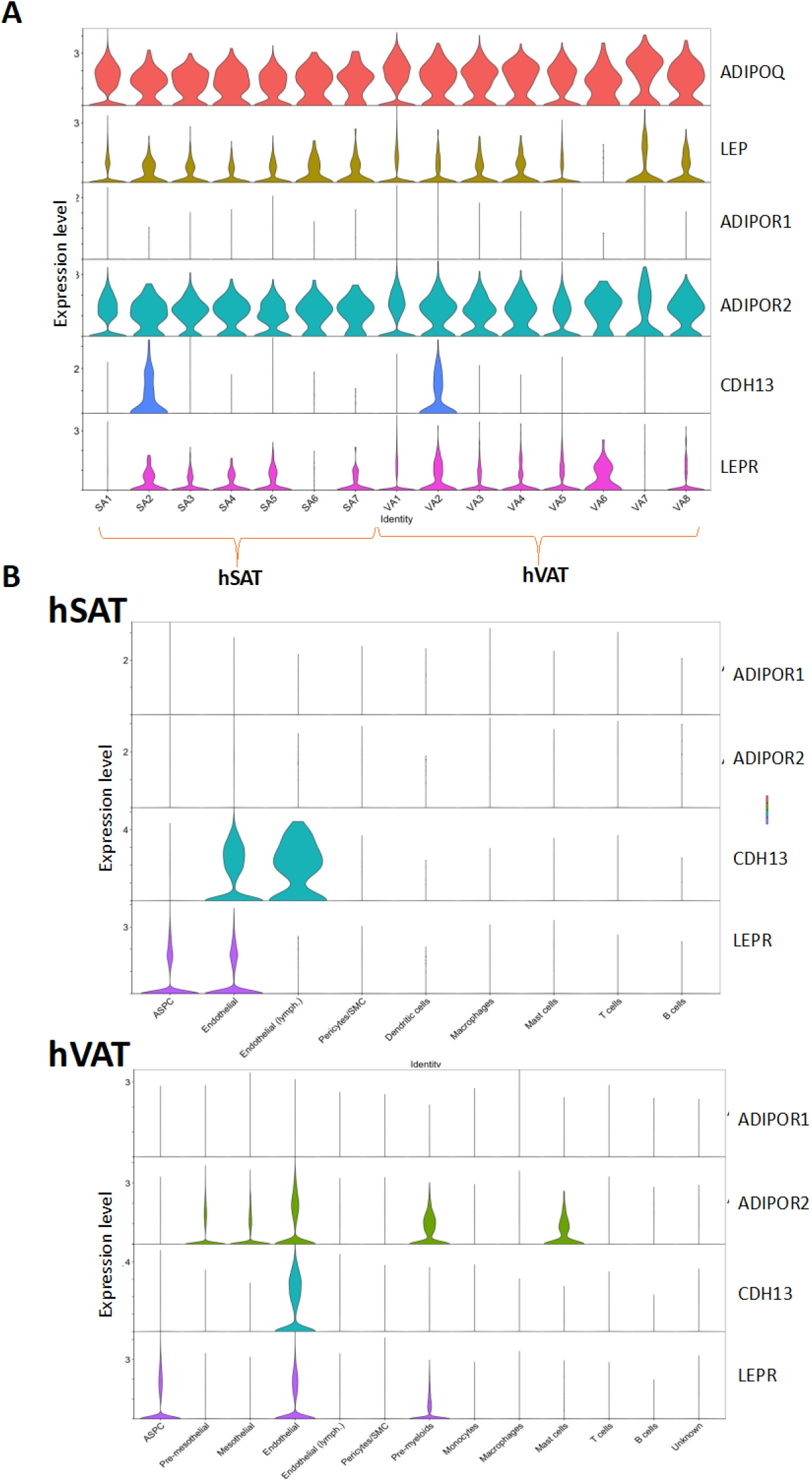
Expression of adiponectin, leptin, and their receptors in hSAT and hVAT. A. Expression levels in adipocyte subtypes of hSAT (left) and hVAT (right). B. Expression levels in non-adipocyte cell-types of each depot.

**Figure S7.**
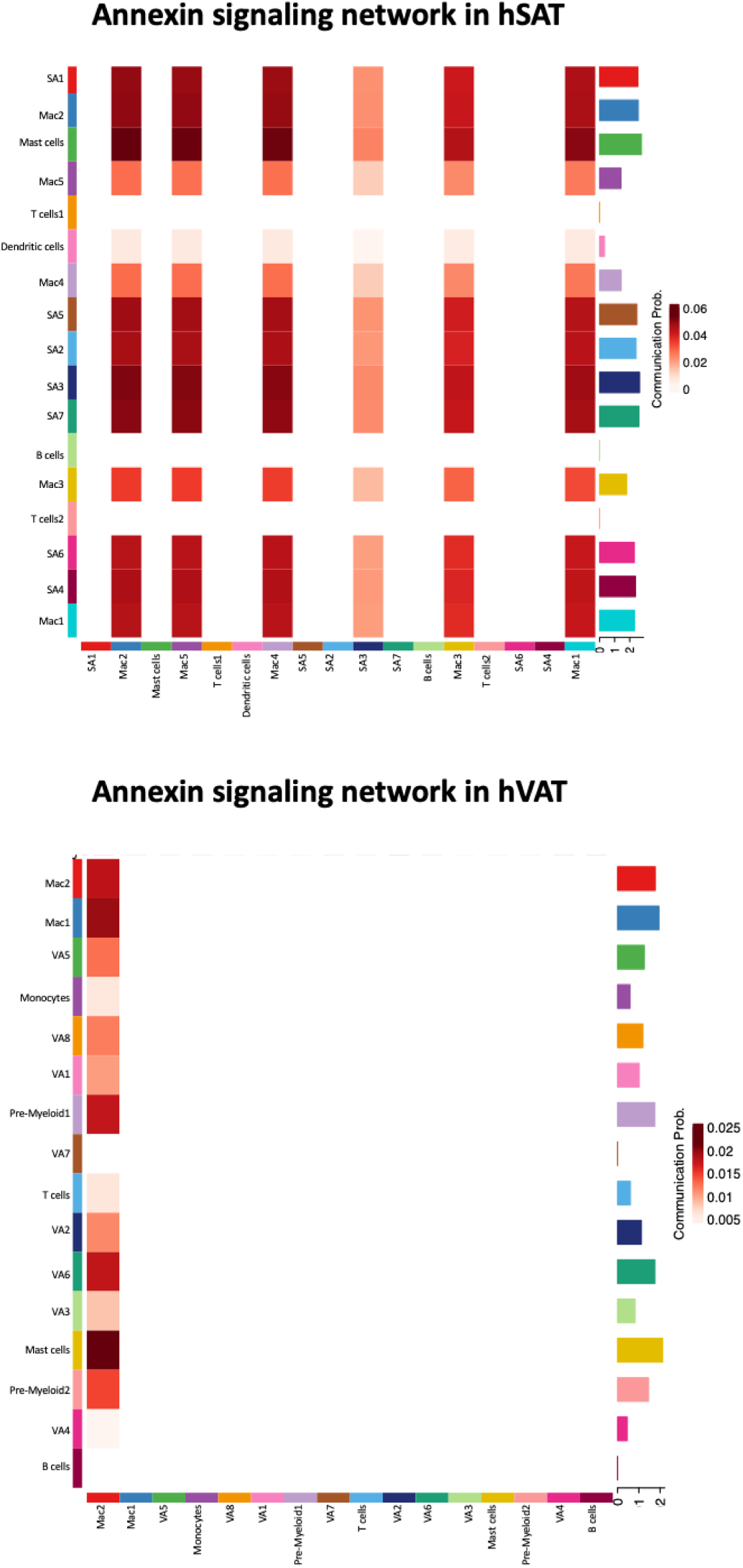
ANNEXIN signaling network in hSAT and hVAT. The ANNEXIN signaling network is much more abundant in hSAT compared to hVAT. In both depots most cells acts as senders. In hSAT all macrophages and immune-related adipocyte subtype (SA3) are targets in hSAT, whereas in hVAT only one macrophage group is targeted. The likelihood of interactions is also higher in hSAT versus hVAT (0.06 compared to 0.025).

**Figure S8.**
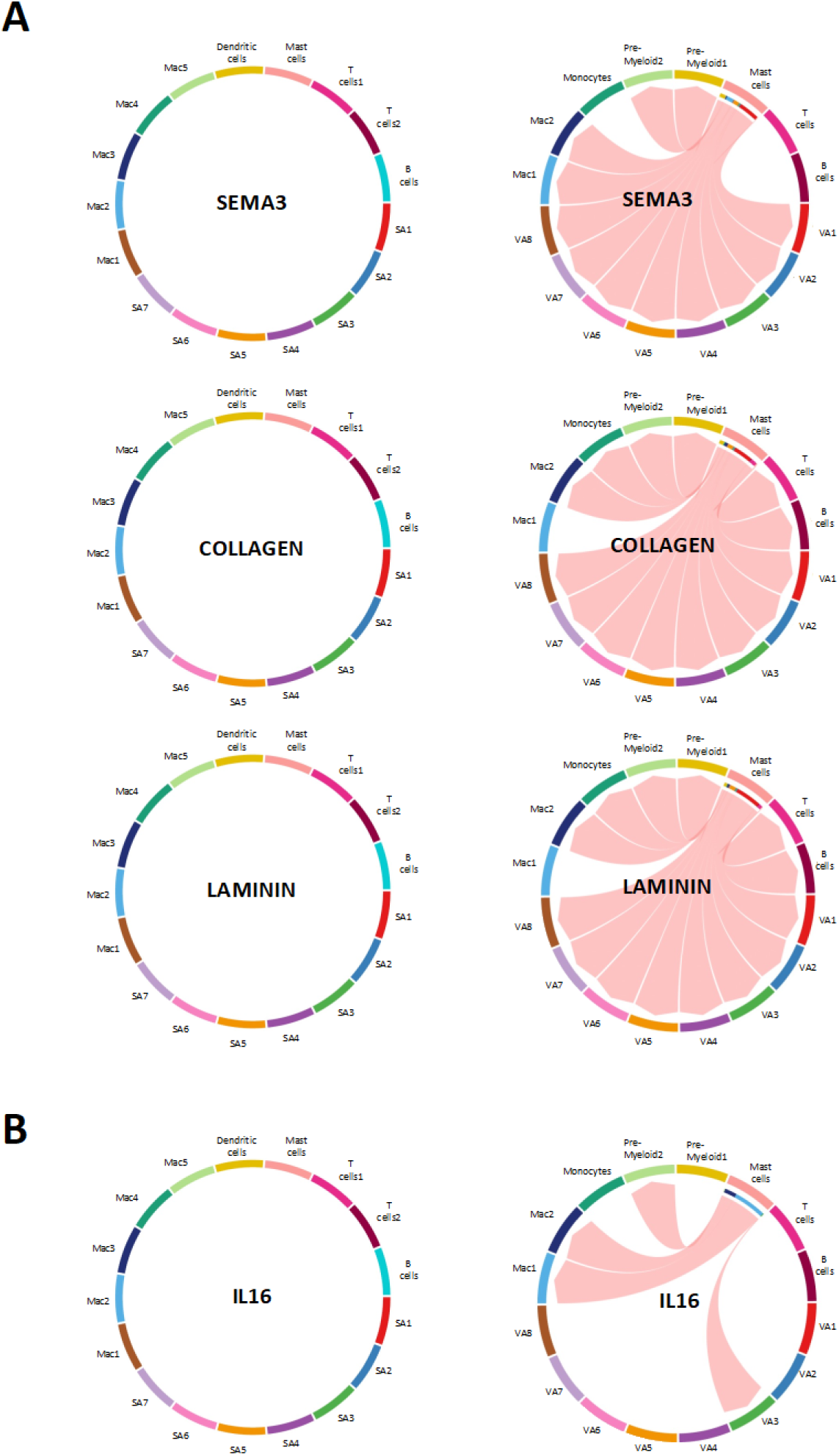
Mast cells involvement as source for ECM, fibrotic and inflammatory pathways differs between depots. A. In hVAT but not in hSAT, mast cells are senders of ECM and fibrotic pathways SEMA3, COLLAGEN and LAMININ. B. In hVAT but not in hSAT, mast cell are senders of the pro-inflammatory (IL16) signaling pathway.

